# Transfreq: a Python package for computing the theta-to-alpha transition frequency from resting state EEG data

**DOI:** 10.1101/2021.12.03.471064

**Authors:** Elisabetta Vallarino, Sara Sommariva, Dario Arnaldi, Francesco Famà, Michele Piana, Flavio Nobili

**Affiliations:** Dipartimento di Matematica (DIMA), Università degli Studi di Genova, Via Dodecaneso 35, 16147, Genoa, Italy; CNR-SPIN, Corso Perrone 24, 16152, Genoa, Italy; Dipartimento di Neuroscienze, Riabilitazione, Oftalmologia, Genetica e Scienze Materno-Infantili (DINOGMI), Università degli Studi di Genova, Largo Daneo 3, 16132, Genoa, Italy; IRCCS Ospedale Policlinico San Martino, Largo Rosanna Benzi 10, 16132, Genoa, Italy

**Keywords:** Transition frequency, Quantitative EEG, Power Spectrum, Clustering, Machine Learning, Neurodegenerative diseases

## Abstract

A classic approach to estimate the individual theta-to-alpha transition frequency requires two electroencephalographic (EEG) recordings, one acquired in restingstate condition and one showing an alpha de-synchronisation due e.g. to task execution. This translates into longer recording sessions that my be cumbersome in studies involving patients. Moreover, incomplete de-synchronisation of the alpha rhythm may compromise the final estimation of the transition frequency. Here we present *transfreq*, a Python library that allows the computation of the transition frequency from resting-state data by clustering the spectral profiles at different EEG channels based on their content in the alpha and theta bands. We first provide an overview of the *transfreq* core algorithm and of the software architecture. Then we demonstrate its feasibility and robustness across different experimental setups on a publicly available EEG data set and on in-house recordings. A detailed documentation of *transfreq* and the codes for reproducing the analysis of the paper with the open-source data set are available online at https://elisabettavallarino.github.io/transfreq/

## 1. Introduction

The analysis of resting state EEG power spectra is a reliable and cheap tool for studying both normal aging (Soininen et al., 1982; Babiloni et al., 2006) and neurodegenerative brain diseases (Malek et al., 2017; Moretti et al., 2004; Klassen et al., 2011). For example, there is evidence that the EEG power in the alpha band and in the slow-wave frequency bands (e.g. theta and delta) shows a direct and an inverse correlation with cognitive performances, respectively. This result has been exploited to support the discrimination of patients affected by the most common neurodegenerative brain diseases from healthy controls (Klimesch, 1999; Jaramillo-Jimenez et al., 2021; Özbek et al., 2021). However, such harmonic behaviors often present significant individual differences (Donoghue et al., 2020) and, moreover, alpha and theta bands, whose power expresses opposite pathophysiological meanings, are contiguous. Therefore, at the individual level the risk is consistent that part of the alpha power band is included in the range of the theta power (i.e., 4-8 Hz), thus implying a wrong interpretation of its (patho)physiological meaning. Establishing the theta-to-alpha transition frequency (TF) at an individual level is therefore of paramount importance in order to avoid misinterpretation of quantitative EEG (qEEG) data. The availability of a computational tool for the determination of TF represents a crucial prerequisite for a meaningful usability of frequency-band power analysis for both research and clinical purposes.

The current standard for TF determination is represented by a more than twenty years old study performed by Klimesch (1999). This approach relies on the fact that event-related de-synchronisation induces a decrease of the alpha power and an increase of the theta power of the event-related power spectrum, with respect to the power spectrum measured during resting state (Klimesch et al., 1997). It immediately follows that theta-to-alpha TF can be determined by comparison between the task-related and the resting state power spectra. The Klimesch’s approach has been successfully used in a number of papers (Singh et al., 2015; Moretti et al., 2004, 2007; Saad et al., 2018). However, its main drawbacks are that (i) it needs the acquisition of two data sets, i.e. a resting state and a event-related time series; and (ii) the task utilised for event-related recording must induce changes in the power spectrum significant enough to allow the identification of variations in the alpha and theta power.

The present study introduces *transfreq*, a publicly-available Python package implementing a novel algorithm for the automated computation of TF from theta to alpha band that works even when just resting-state EEG time series are available. This computational approach relies on the determination of appropriate features associated to the power spectrum measured at each channel, and on the application of an unsupervised algorithm that automatically identifies two clusters of EEG sensors associated to the alpha and theta bands, respectively. In *transfreq* we implemented four different strategies for selecting the sensor-level features and the corresponding clustering algorithms (Saxena et al., 2017). The workflow of these approaches is illustrated in the case of a test-bed example and validated on both an open-source data set and time series recorded during an experiment performed in our lab. For most subjects in both data sets *transfreq* estimate a value of TF close to that obtained by using the Klimesch’s method. Additionally, we show some typical scenarios in which the classic Klimesch’s method fails in capturing the correct TF while *transfreq* still returns plausible estimates.

## 2. Materials and methods

### 2.1. Klimesch’s method

A classic approach to compute theta-to-alpha TF is that proposed by Klimesch and colleagues (Klimesch, 1999) and schematically depicted in Figure 1A. In detail, the Kilmesch’s method requires two EEG recordings as input, one acquired during a resting-state condition and one acquired while the subject is performing a task. For both recordings and for each one of the *N* EEG sensors, the power spectrum (Vallarino et al., 2020; Bendat and Piersol, 2011) of the corresponding time series is computed and normalised by dividing for the norm over all frequencies, i.e., we computed

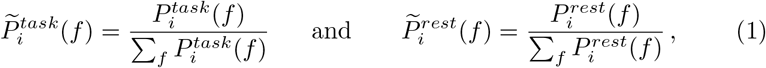

where 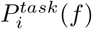 and 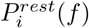 are the power spectra at frequency *f* of the signal recorded by the *i*-th sensor during the task and the resting–state conditions, respectively. Then, the mean over all the EEG channels of the normalised power spectra in (1) is computed to obtain two spectral profiles, namely

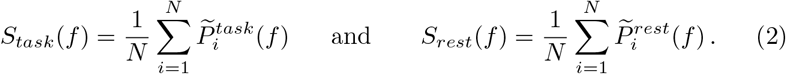

**Figure 1:**
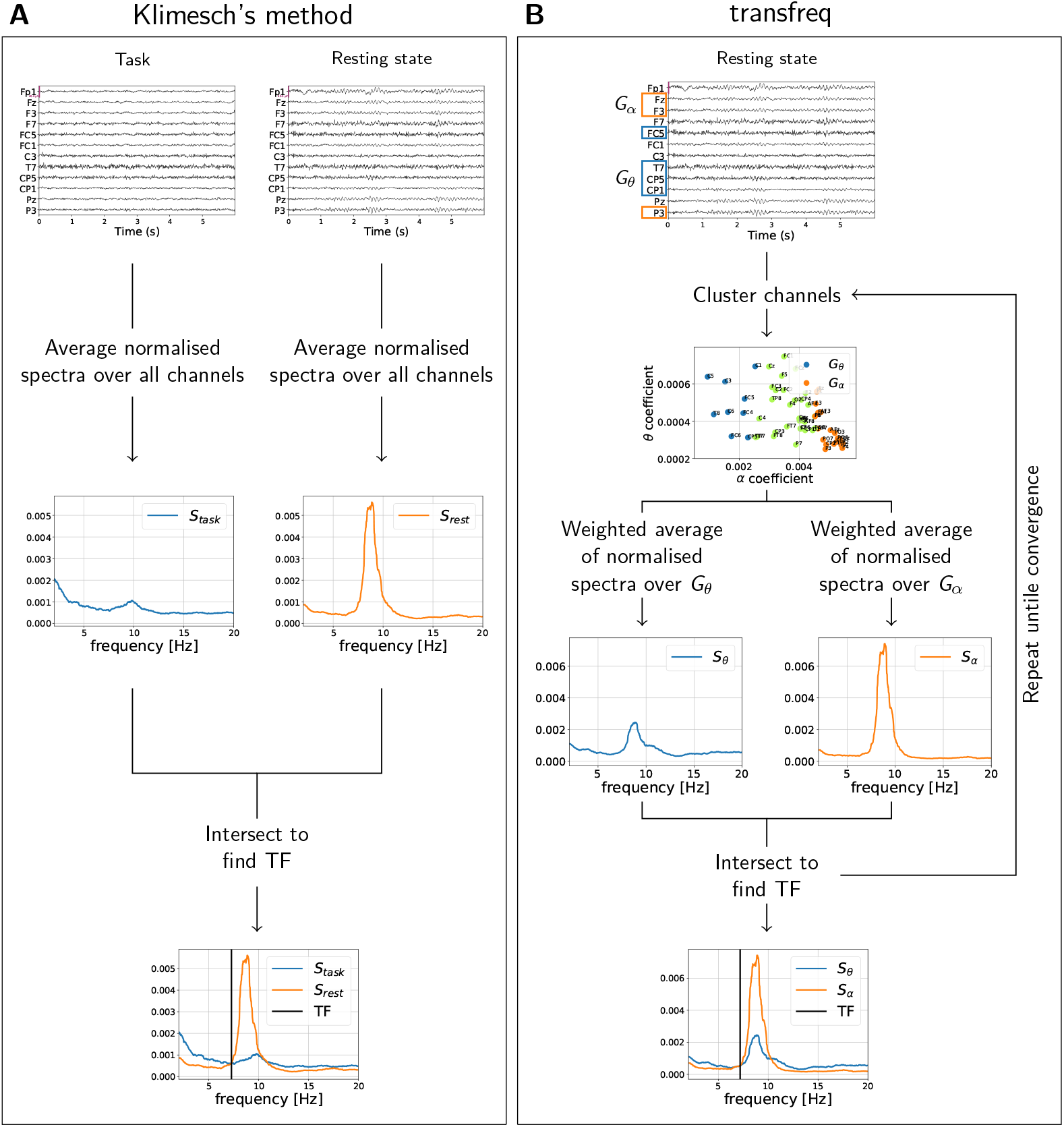
Comparison between the pipelines of the classic Kilmesch’s method (A) and of *transfreq* (B).

The Klimesch’s method relies on the fact that *S_rest_* usually presents a peak in the alpha band while, due to task-related alpha de-synchronisation, *S_task_* presents a lower intensity in the alpha band and a higher intensity in the theta band with respect to *S_rest_* (Klimesch, 1996; Klimesch et al., 1998; Schacter, 1977). TF is thus defined as the highest frequency before the individual alpha peak (IAP) at which *S_task_* and *S_rest_* intersect. Here, the IAP is defined as the frequency in the range [7, 13] Hz at which *S_rest_* peaks (Babiloni et al., 2004).

### 2.2. Transfreq algorithm

In this paper we introduce *transfreq*, a method to automatically compute the TF from theta to alpha band when only resting-state EEG data are available. *Transfreq* relies on a rationale similar to that of Klimesch’s method. Namely, TF is defined as the intersection between two spectral profiles differing in their content within the alpha and theta bands. However, with respect to the Klimesch’s method, such profiles are computed by exploiting the fact that alpha and theta activities are not uniformly expressed across the different EEG channels. In fact, some channels present high alpha activity (typically, channels above the occipital lobe), whereas others show lower alpha and higher theta activities (typically, channels corresponding to temporal and frontal brain areas) (Klimesch, 1996; Nunez et al., 2001). Consequently, two groups of channels can be identified: the first group includes channels characterized by a preponderant alpha activity (this group plays a role analogous to the one of EEG data measured at rest in Klimesch’s method); the second group includes channels showing preponderant theta activity and limited alpha activity (this second group plays a role analogous to the one of the task-evoked EEG recordings in Klimesch’s method).

The *transfreq* pipeline is schematically illustrated by Figure 1B and Algorithm 1. In detail, for each EEG channel the normalised power spectrum is computed as in equation (1), that is

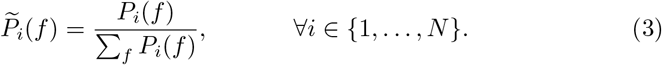

TF is determined through the following iterative procedure.

i. Set an initial value for the alpha and theta frequency-bands. Specifically, the alpha frequency-band is identified as a 2 Hz range centred on the IAP, which is defined as the frequency where the power spectrum averaged over all sensors peaks; the theta frequency-band is set equal to [5, 7] Hz, or to [IAP – 3, IAP – 1] Hz if the previous interval overlaps with the alpha frequency-band.
ii. Compute, for each channel, the alpha and theta coefficients by averaging the normalised power spectrum 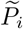 over the corresponding frequency band.
iii. Apply a clustering algorithm to identify two groups of channels based on the alpha and theta coefficients. The channels in the first group, denoted as *G_θ_*, will be characterised by low alpha and high theta activities, while the channels in the second group, *G_α_*, will be characterised by high alpha and low theta activities. Two spectral profiles are thus obtained through a weighted average of the power spectra over the two groups, that is

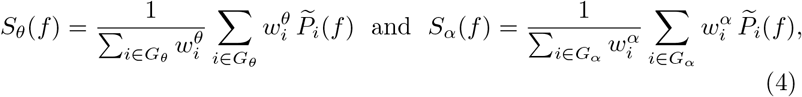

where 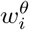 and 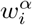 are the theta and alpha coefficients for channel *i*, respectively.
iv. Define a first estimate of TF as the highest frequency before the IAP at which *S_θ_* and *S_α_* intersect.
v. Use the value of TF computed in (iv) to define new, more accurate, alpha and theta frequency bands, set equal to [max{*IAP* – 1,*TF*}, *IAP* + 1] and [*TF* – 3, *TF* – 1], respectively. Such a choice guarantees the intervals to be fully characterised by alpha and theta activation. Indeed, we chose narrower bands with respect to the classic 4 Hz ranges defined in the literature (Bazanova and Vernon, 2014; Klimesch, 1999) and we impose at least a 1 Hz separation between the intervals.

Steps (ii)-(v) are iterated until a desired level of accuracy is reached, quantified as the difference between two consecutive estimates of TF. The desired level of accuracy is set equal to the highest value between 0.1 Hz and the frequency resolution Δ_*f*_. The rationale behind this choice is that 0.1 Hz is an acceptable error when computing TF. However, if the frequency resolution is lower (i.e Δ_*f*_ > 0.1 Hz), setting the desired level of accuracy to 0.1 Hz would be the same as setting it to 0, which is a too strong requirement; therefore in such cases the level of accuracy is set equal to the frequency resolution.

We point out that the effectiveness of *transfreq* depends on the clustering procedure used to define the two groups of channels *G_θ_* and *G_α_*. In *transfreq* we have implemented four different algorithms, described in the next sub-sections.

#### 2.2.1. Clustering method 1: 1D thresholding

The first clustering method implemented in *transfreq* is based on the ratio between the alpha and theta coefficients computed for each channel. In fact, channels with a low value of such alpha-to-theta ratio are characterised by low alpha and high theta activities, whereas channels with a high value are characterised by high alpha and low theta activities. The first group of channels, *G_θ_*, is thus defined by the four channels showing the lowest values of the alpha-to-theta ratio, while the second group, *G_α_*, is defined by the four channels showing the highest values of the same ratio. A visual representation of this approach on a representative data set can be seen in Figure 2A. In *transfreq*, the number of channels in each group has been set equal to 4 after computing and visually inspecting the results for different values of such a parameter. In fact, the overall behaviour of the algorithm was similar across the different tested values.

**Algorithm 1.**
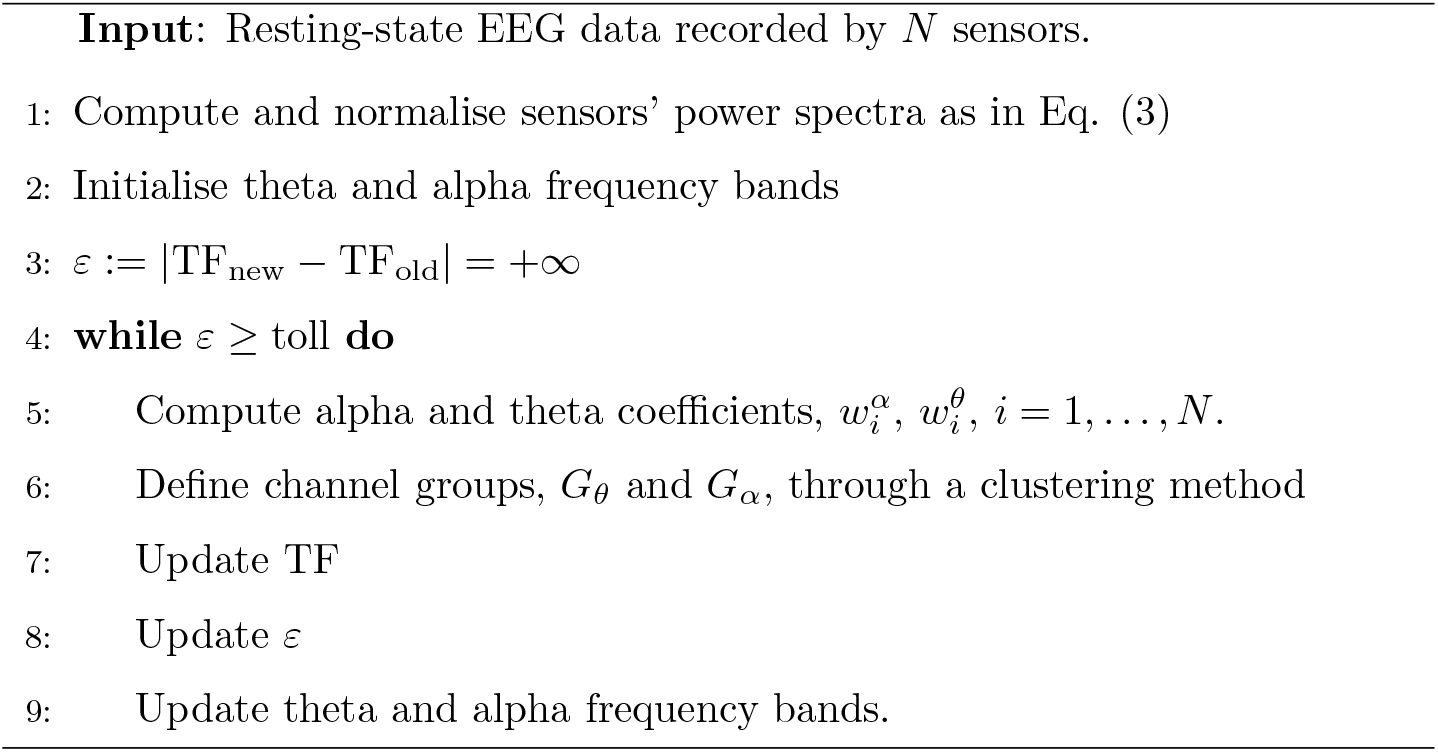
*transfreq* core algorithm

**Figure 2:**
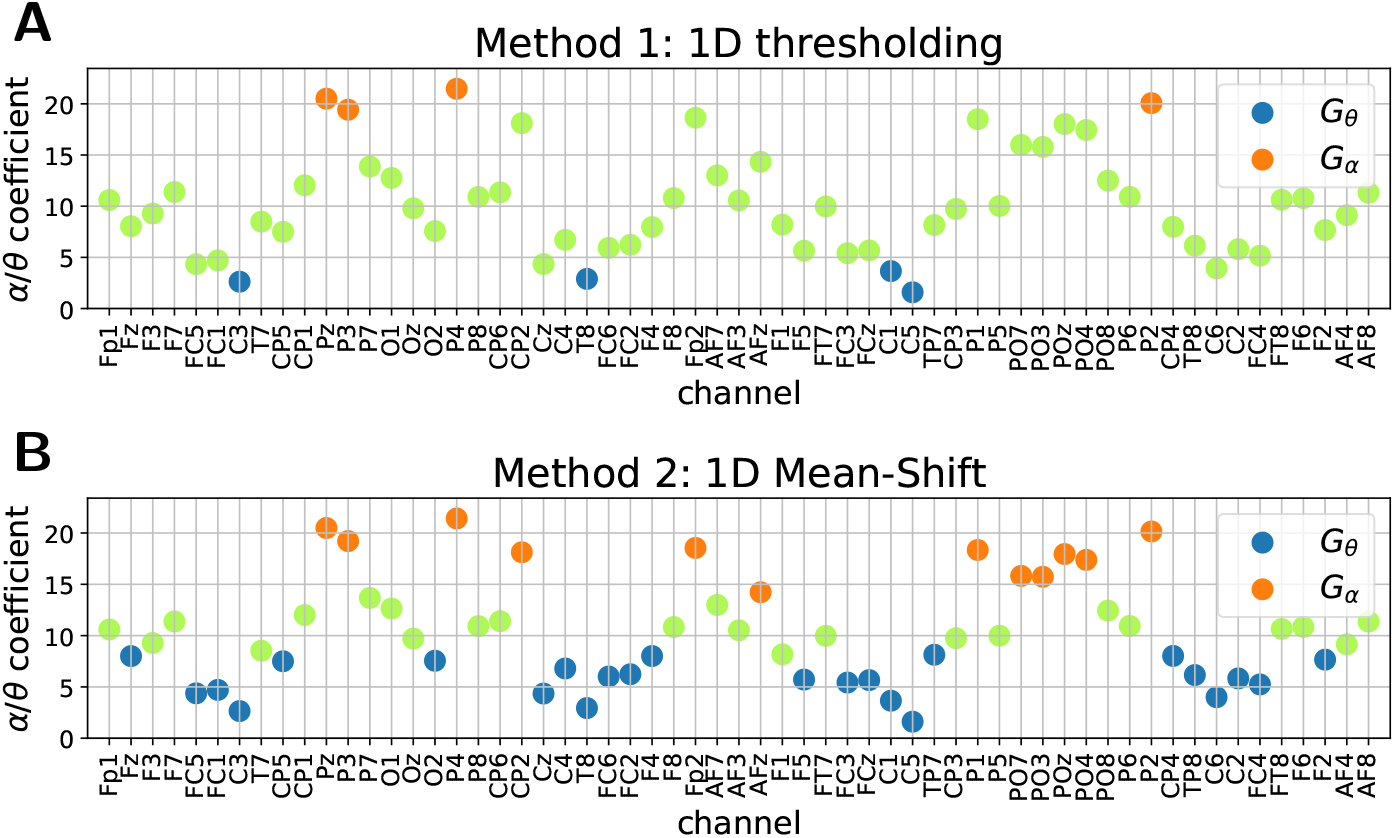
Performance illustration of the 1D clustering approaches thresholding (A) and mean-shift (B). Both panels show the value of the ratio between alpha and theta coefficients as function of the EEG sensors. Channels that belong to *G_θ_* and *G_α_* are represented as blue and orange dots, respectively. In *transfreq*, the remaining channels (greed dots) are excluded from the subsequent analysis.

#### 2.2.2. Clustering method 2: 1D mean-shift

One drawback of the previous approach is the need to heuristically set the number of channels within the two groups *G_α_* and *G_θ_*. To overcome such a limitation, we implemented a second clustering approach where the Mean Shift algorithm (Comaniciu and Meer, 2002) is used to cluster the EEG sensors with respect to the ratio between the alpha and theta coefficients computed, for each channel, as described in the previous sub-section. To this end we used the MeanShift function available within the Python package Scikit Learn (Pedregosa et al., 2011) that also automatically determines the number of clusters. *G_θ_* is then defined equal to the cluster containing the channel with the lowest value of the alpha-to-theta ratio, while *G_α_* is set equal to the cluster containing the channel with the highest value of the same ratio. A visual representation of this approach on a representative data set can be seen in Figure 2B.

#### 2.2.3. Clustering method 3: 2D k-means

Both approaches described in the previous sub-sections rely on 1-dimensional clustering techniques that use the ratio between the alpha and theta coefficients as feature. In the third approach implemented in *transfreq* we exploited the k-means algorithm (Lloyd, 1982) to cluster the EEG sensors by using the alpha and theta coefficients as two distinct features. To this end, we used the KMeans function within the Python package Scikit Learn (Pedregosa et al., 2011). The number of clusters to generate is set equal to 2. Then *G_α_* is defined as the cluster whose centroid shows the highest value of the alpha coefficient, while the other cluster defines *G_θ_*. As illustrated in Figure 3A, channels belonging to *G_α_* (orange dots) typically present a higher alpha coefficient and a lower theta coefficient than the other ones (blue dots).

**Figure 3:**
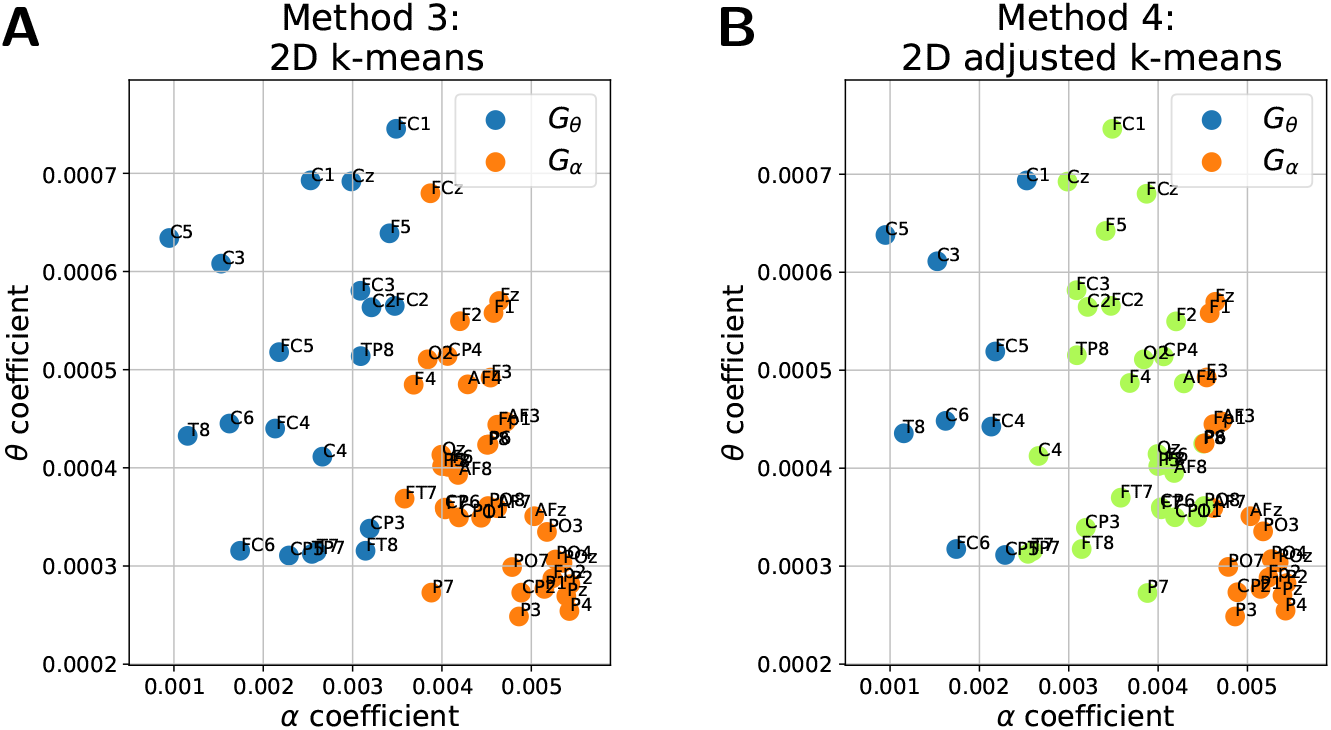
Performance illustration of the 2D clustering approaches k-means (A) and adjusted k-means (B). Both panels show the value of the theta coefficients on the y-axis and that of the alpha coefficients on the x-axis. Channels that belong to *G_θ_* and *G_α_* are represented as blue and orange dots, respectively. In *transfreq*, the remaining channels (green dots) are excluded from the subsequent analysis.

#### 2.2.4. Clustering method 4: 2D adjusted k-means

The fourth clustering approach implemented in *transfreq* takes as input the two sensors groups, *G_α_* and *G_θ_*, computed using the k-means algorithm as described in the previous sub-section. However, the two groups are now adjusted so that only sensors showing the highest inter-cluster difference in terms of the alpha and theta coefficient values are retained. To this end, as illustrated in Figure 3B, we removed from *G_α_* and *G_θ_* all points laying between the two lines that pass through the centroids and are perpendicular to the segment connecting the two centroids.

### 2.3. Software architecture

The approach described in the previous section is implemented in the publicly available Python library *transfreq* (https://elisabettavallarino.github.io/transfreq/). As shown in Table 1, *transfreq* comprises two modules: a set of three operative functions, that allow the estimation of TF either with the classic Klimesch’s method or with our approach, and a set of six functions to visualise the results.

**Table 1:**
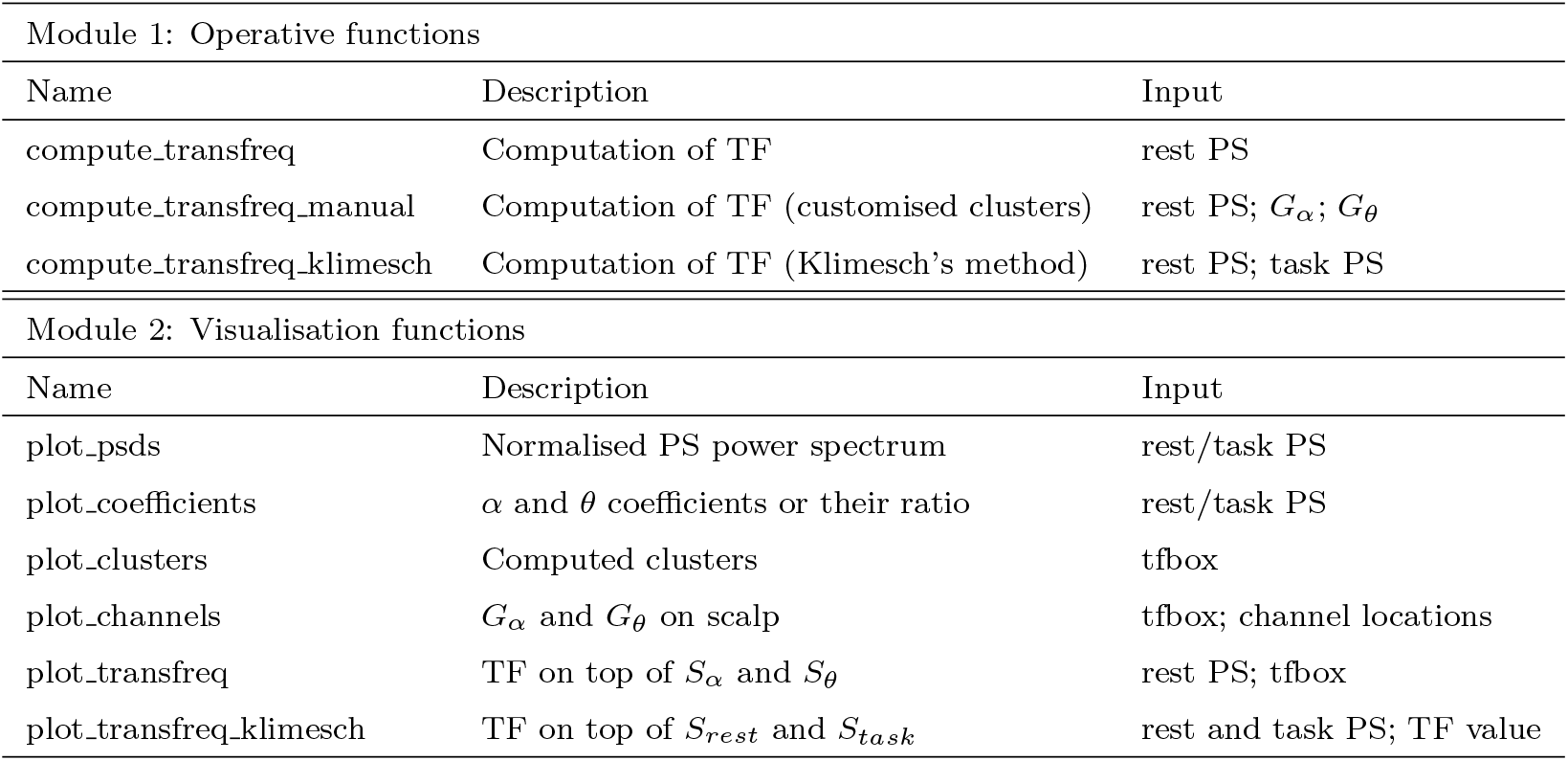
Functions implemented within *transfreq.* The table provides the name of each function (first column), a short description of their purpose (second column), and the required input variables (third column). Here, rest PS and task PS stand for resting state and task-related EEG power spectrum, respectively; tfbox is a dedicated dictionary output of the operative functions. For some of the functions, an additional set of optional arguments may be passed by the user, such as predefined alpha and theta frequency-band, or the clustering approach to be used for defining *G_α_* and *G_θ_*. The full list of these additional parameters may be found in the package documentation.

#### 2.3.1. Module 1: operative functions

All the operative functions require in input the power spectra of the recorded EEG data. These power spectra have to be provided as matrices of size *N* × *F*, where *N* is the number of EEG sensors and *F* is the number of frequencies in which the power spectra are evaluated.

The function *compute_transfreq* implements the iterative procedure described in Algorithm 1. Customised estimation of the transition frequency may be obtained through the function *computeJmnsfreq_manual* by providing two predefined groups of channels *G_α_* and *G_θ_*. In this case, TF is computed by looking at the intersections between the corresponding spectral profiles *S_α_* and *S_θ_*. Both functions return a dedicated dictionary, called *tfbox* in Table 1, that contains: (i) the results of the clustering procedure, together with the alpha and theta coefficients, 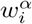 and 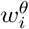, associated to each one of the sensors; (ii) the name of the employed algorithm; (iii) the estimated value of TF.

In order to provide an exhaustive toolbox for computing the theta-to-alpha TF we also implemented a function for the computation of TF with Klimesch’s method. Such a function is named *compute_transfreq_Klimesch* and only returns the estimated value of TF.

#### 2.3.2. Module 2: visualisation functions

As shown in Table 1, *transfreq* offers the users two functions to visualise features of the data provided in input, namely the normalised EEG power spectrum (function *plot_psds*) and the corresponding alpha and theta coefficients (function *plot_coefficients*).

Three other functions allow the user to visualise results from each step of our approach, that is: (i) the alpha and theta coefficients grouped according to the results of the clustering procedure (function *plot_clusters*); (ii) the corresponding channels group *G_α_* and *G_θ_* located on top of topographical maps (function *plot_channels*); (iii) the final estimated value of TF on top of the spectral profiles *S_α_* and *S_θ_* (function *plot_transfreq*). The function *plot_channels* makes use of the Python package visbrain (Combrisson et al., 2019), and, in particular, we modified its function *TopoObj* to optimise it to our visualisation purpose.

Eventually, the function *plot_transfreq_klimesch* is dedicated to plot the value of TF estimated using the classic Klimesch’s method.

### 2.4. Data

We validated *transfreq* by using two EEG data sets. The first one is an open-source data set, while the second one is an in-house data set we recorded in our lab. We used two different data sets to test the robustness of *transfreq* across data recorded in different experimental conditions.

#### 2.4.1. Open-source data set

This data set contains EEG data available at OpenNeuro, a free and open platform for sharing neurophysiological data (Gorgolewski et al., 2017), at the accession number ds003490 (data set DOI doi:10.18112/openneuro.ds003490.v1.1.0). Data comprise both resting state and stimulus auditory oddball EEG recordings, sampled at 500 Hz, from 25 Parkinson’s patients and 25 matched controls. For Parkinson’s patients, two sessions are available, while for healthy controls one session is available. More information about this data set can be found in the paper by Cavanagh et al. (2018). For each subject and for each session we selected two minutes of recording under stimulation, and one minute resting state eyes-closed recording.

#### 2.4.2. In-house data set for validation

This data set included 80 traces acquired during a previous multicenter study, namely the Innovative Medicines Initiative PharmaCog project: a European ADNI study (Galluzzi et al., 2016). This study aimed at investigating multiple biomarkers in a population with amnesic mild cognitive impairment (MCI), by following subjects for three years or until conversion to dementia. EEG was repeatedly acquired every 6 months; thus the 80 traces refer to 16 subjects undergoing EEG from one to 7 times. The 16 subjects (8 males, 8 females, age range 55-82 years, mean: 70±6 years; mini-mental state examination score range at first evaluation: 23-30, mean: 26.5±2.13) included 11 who converted to Alzheimer disease dementia during the follow-up, 2 subjects who convert to frontotemporal dementia, and 3 subjects who remained in an MCI stage or even reverted to a normal condition.

For the analysis we selected two and a half minutes of resting state eyes-closed recording and two and a half minutes of resting-state eyes-opened recording, where data showed a de-synchronisation of the alpha rhythm (Gómez-Ramírez et al., 2017). Both data were recorded with a sampling frequency of 512 Hz.

### 2.5. Data analysis

The recorded time series from both data sets were first pre-processed using the MNE-Python analysis package (Gramfort et al., 2013). For each subject and for each condition, the EEG recording was filtered between 2 and 50 Hz, while bad segments were manually removed and bad channels were interpolated. Then, data were re-referenced using average reference (Offner, 1950) and Indepentend Component Analysis (ICA) (Jutten and Herault, 1991) was applied for artefact and noise removal. Remaining bad segments were automatically rejected by using the autoreject Python package (Jas et al., 2017). Finally, the pre-processed EEG recordings were visually inspected by experts and discarded when they did not present a visible alpha peak. In this way, in the open-source data set we excluded the first session of four subjects and both sessions of one subject. In the in-house data set all sessions involving four subjects were excluded from the analysis.

Power spectra were computed in the 2-30 Hz range with the multitapers method (Thomson, 1982). With such a method the frequency resolution of the power spectra depends on the time resolution and duration of the EEG recordings. In order to apply the Klimesch’s method, the spectral profiles under the two conditions (rest and task) need to have the same frequency resolution. To this end the length of both recordings was set equal to the length of the shortest one. Average duration of the EEG recordings from the open source data set was 58 s, while average duration of the EEG recordings from the in-house data set was 134 s. Afterwards, TF was computed using both the Klimesch’s method and *transfreq.* Finally, the results obtained with Klimesch’s method were visually inspected by experts and excluded when the method did not provide reliable results. Exclusion criteria comprised cases in which the two spectral profiles did not intersect as well as cases in which the two spectral profiles overlapped. This process led to the exclusion of 19 EEG recordings from the open-source data set and 14 EEG recordings from the in-house data set. Therefore, the analysis to validate *transfreq* was performed on a total of 50 EEG recordings from the open-source data set and 45 from the in-house data set.

## 3. Results

### 3.1. Transfreq performances on an illustrative example

We first tested the performances of *transfreq* when applied to an illustrative example picked up from the open-source database. Figure 4 and Figure 5 show the results provided by the tool when the four different clustering algorithms were applied. For all algorithms, the resulting *G_α_* mainly contained channels that lie over the occipital lobe and showed a higher alpha activity than the channels in *G_θ_*.

**Figure 4:**
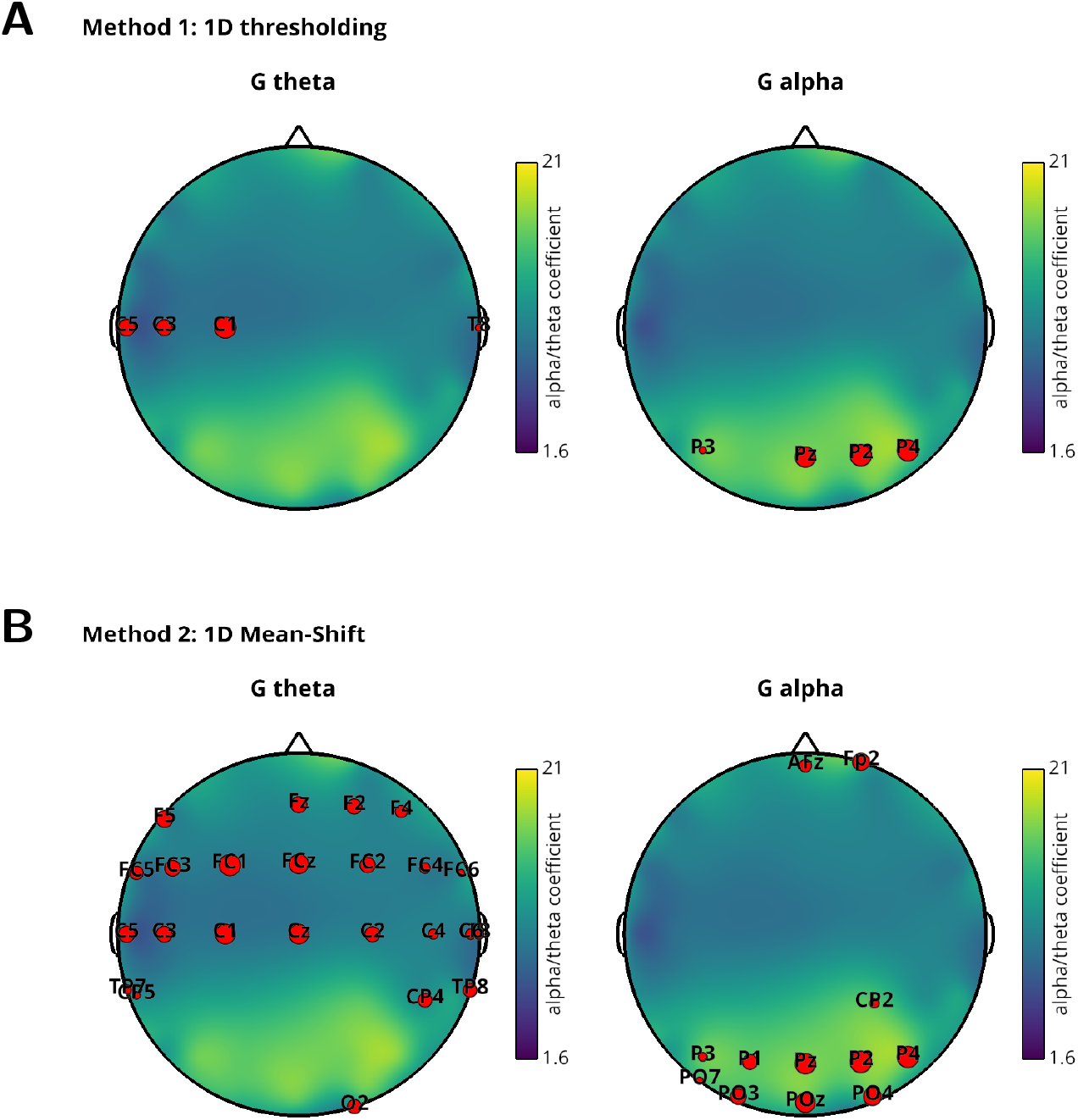
Location on the scalp of channels in *G_θ_* (left column) and *G_α_* (right column) for one representative subject from the open-source data set. Sensors have been clustered by using 1D thresholding (upper row) or 1D mean-shift (lower row). In each panel, red dots represent the selected channels and, in the background, the topographical map shows the value of the ratio between alpha and theta coefficients. For the sensors in *G_θ_*, the size of the dots is proportional to the corresponding theta coefficient, 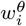, while for those in *G_α_* the size is proportional to the alpha coefficient, 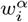.

**Figure 5:**
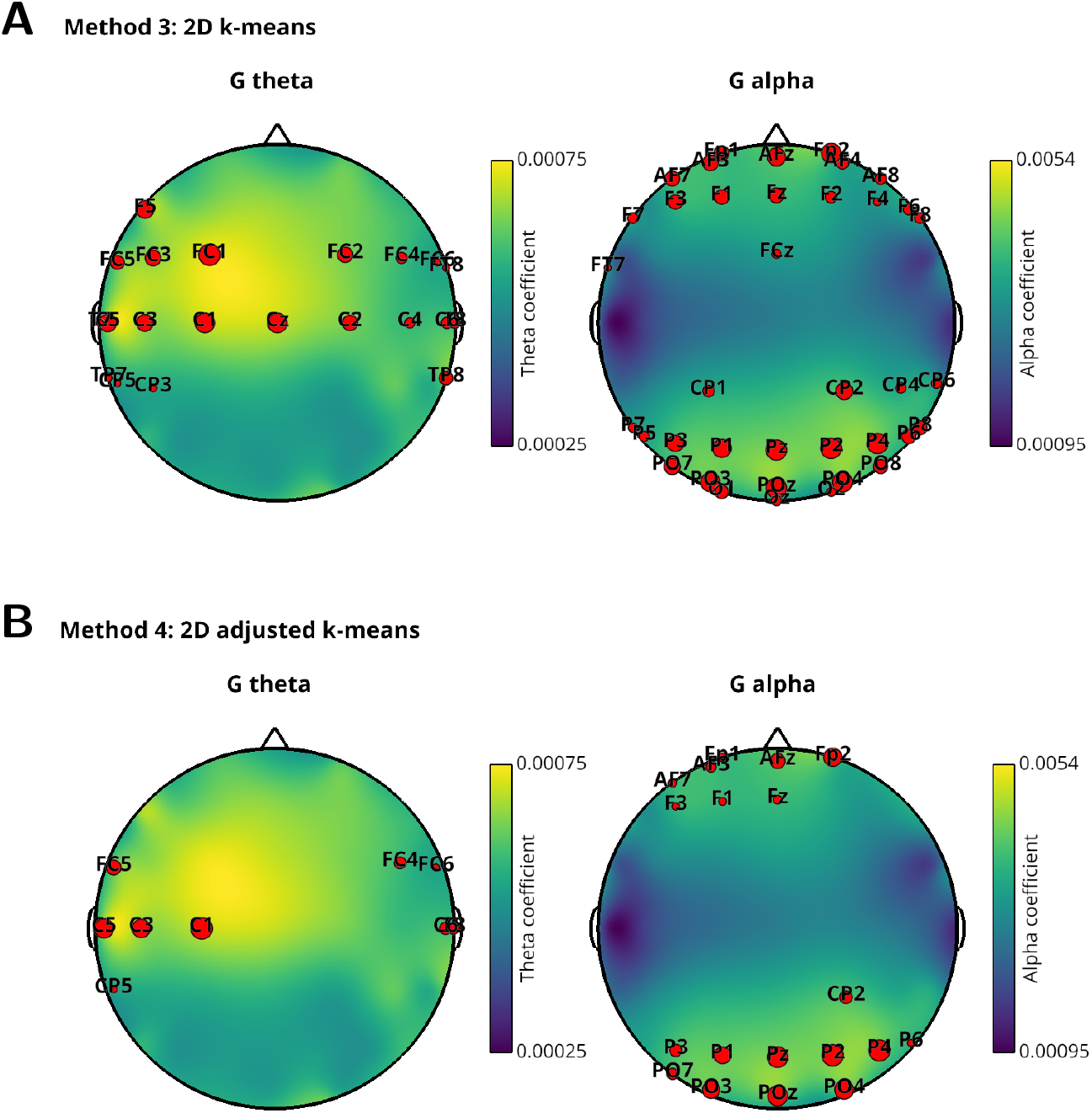
Location on the scalp of channels in *G_θ_* (left column) and *G_α_* (right column) for one representative subject from the open-source data set. Sensors have been clustered by using 2D k-means (upper row) or 2D adjusted k-means (lower row). As in Figure 4, the red dots depict the selected channels. In the two panels on the left side, referring to *G_θ_*, the size of the sensors and the background topographical maps represent the theta coefficient, 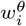. Instead, the two panels on the right side, referring to, *G_α_*, show the alpha coefficients 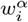.

When 1D thresholding is used, both *G_α_* and *G_θ_* contain a pre-defined number of sensors (4 in this case). Instead, the other methods automatically estimate the size of *G_θ_* and *G_α_*, and thus the two groups may contain a different number of channels.

While with the 2D k-means *G_α_* and *G_θ_* span all the EEG channels, the 2D adjusted k-means starts from the two groups defined by using k-means and selects only the channels showing a high inter-cluster difference. Specifically, as illustrated in Figure 5, the channels in *G_α_* (*G_θ_*) showed both a high alpha (theta) activity and a low theta (alpha) activity.

Depending on the selected clustering approach, *transfreq* may return different estimates for TF, as illustrated in Figure 6. With this subject, the value of TF estimated by the Klimesch’s method was 7.29 Hz, while transfreq returned 7.38 Hz with 1D thresholding, 7.39 Hz with 1D mean-shift, 7.22 Hz with 2D k-means, and 7.19 Hz with 2D adjusted k-means.

**Figure 6:**
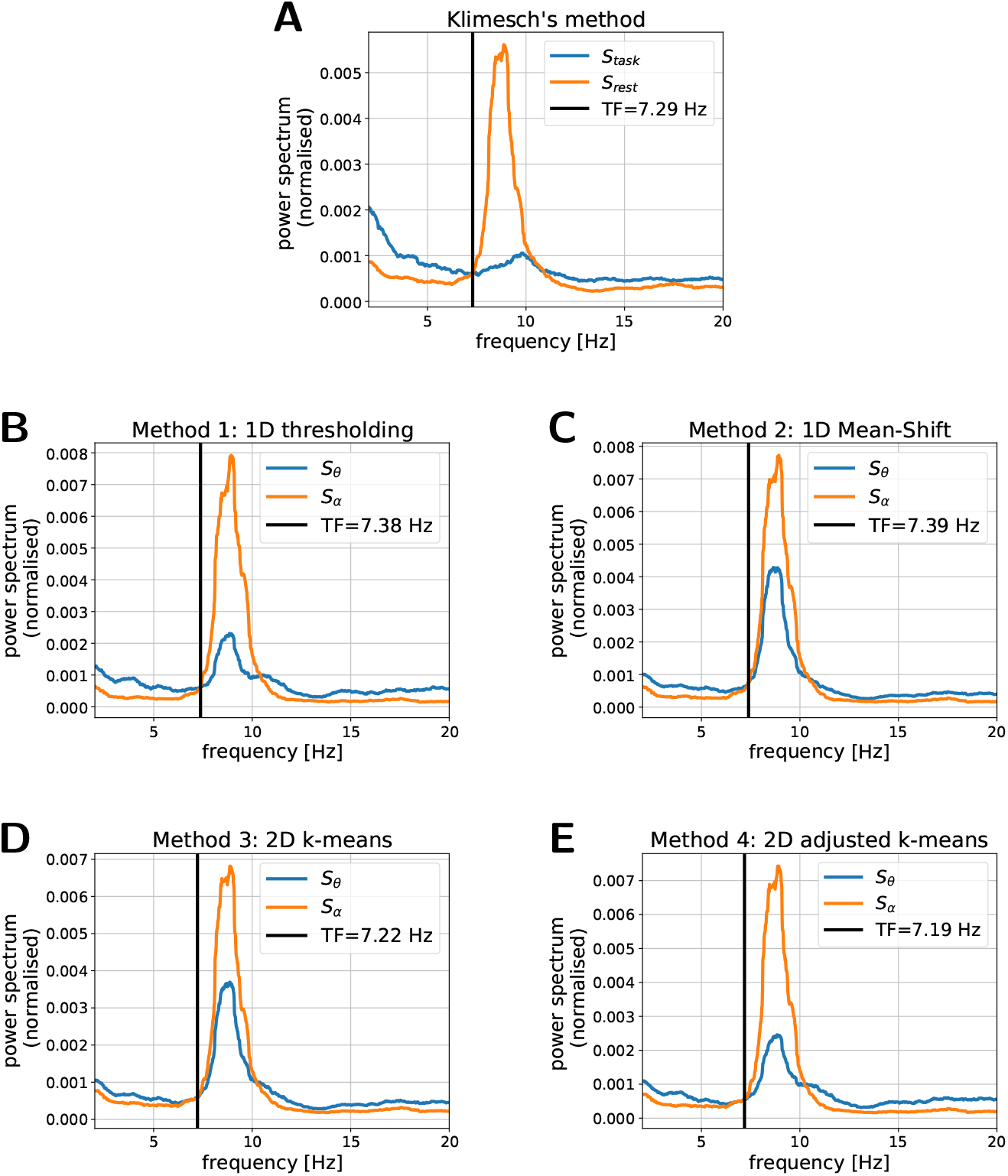
TFs estimated with Klimesch’s method and with *transfreq* by means of the four clustering methods for one representative subject from the open-source data set. In each panel: the blue line depicts the spectral profile with low alpha and high theta activation, namely *S_task_* in Klimesch’s method and *S_θ_* in *transfreq*; the orange line shows the spectral profile with high alpha and low theta activation, namely *S_rest_* in Klimesch’s method and *S_α_* in *transfreq;* the red vertical line indicates the estimated value of TF.

### 3.2. Validation on the open-source data set

As illustrated in Figure 7, for most subjects in the open-source data set, the difference Δ_*TF*_ between the TF value estimated by *transfreq* and by Klimesch’s method was in absolute value below 1 Hz. Specifically, |Δ_*TF*_ | was lower than 1 Hz for 82% of the subjects when 1D thresholding was employed for clustering, 76% in the case of 1D mean-shift, 82% for 2D k-means, and 88% for 2D adjusted k-means. Figure 7 also shows that *transfreq* mainly estimated a lower value of TF than Klimesch’s method. Since the lowest values of |Δ_*TF*_| were obtained by clustering the EEG channels by means of the 2D adjusted k-means method, this is suggested as the default approach within *transfreq*.

**Figure 7:**
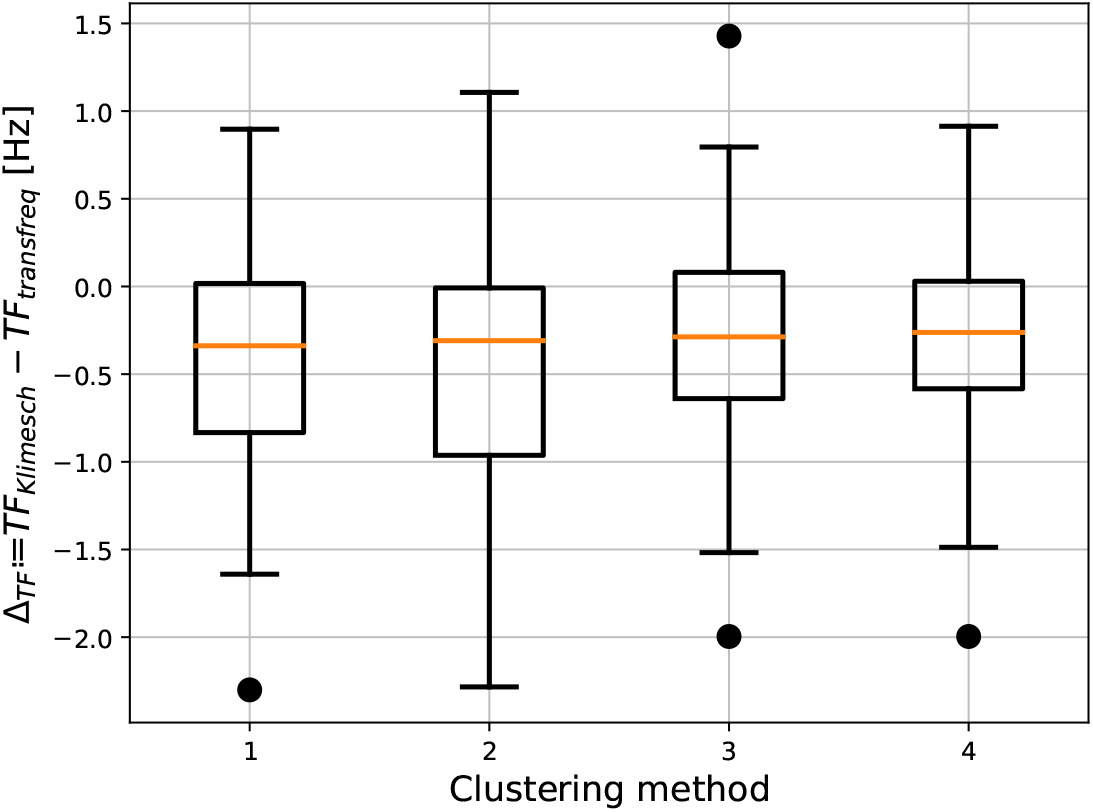
Difference between TFs estimated with Klimesch’s method (*TF_Klimesch_*) and with *transfreq* (*TF_transfreq_*) over the open-source data set. Each boxplot depicts the results obtained when a different clustering approach is used to define the channels group *G_θ_* and *G_α_*, namely: 1D thresholding (Method 1); 1D mean-shift (Method 2); 2D k-means (Method 3); and 2D adjusted k-means (Method 4).

### 3.3. Improvements of transfreq over the Klimesch’s method

Klimesch’s method relies on an event-related reduction of the alpha activity that may not occur in practical scenarios due, for example, to an incorrect execution of the task. Indeed, as shown in Figure 8, for some of the subjects in the considered data sets the spectral profiles *S_task_* and *S_rest_* perfectly overlapped and thus Klimesch’s method failed in computing TF.

**Figure 8:**
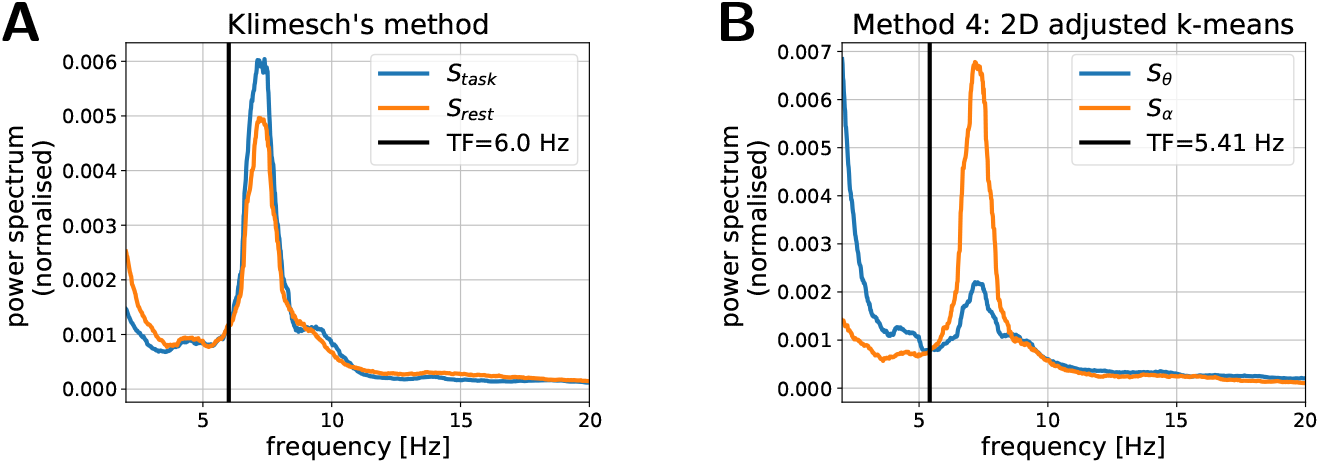
Example where Klimesch’s method provides unreliable estimate of TF because event-related, *S_task_*, and resting-state, *S_rest_*, spectral profiles overlap. (A) Results obtained with the Klimesch’s method. (B) Results obtained with *transfreq* by using 2D adjusted k-means to compute the spectral profiles *S_θ_* and *S_α_*

On the other hand, some subjects may show an event-related modulation of the alpha frequency (Haegens et al., 2014). As represented in Figure 9, in this case the shift of the alpha peak in *S_task_* prevented the use of Klimesch’s method because the two spectral profiles *S_task_* and *S_rest_* did not intersect in the range [0, 10] Hz.

**Figure 9:**
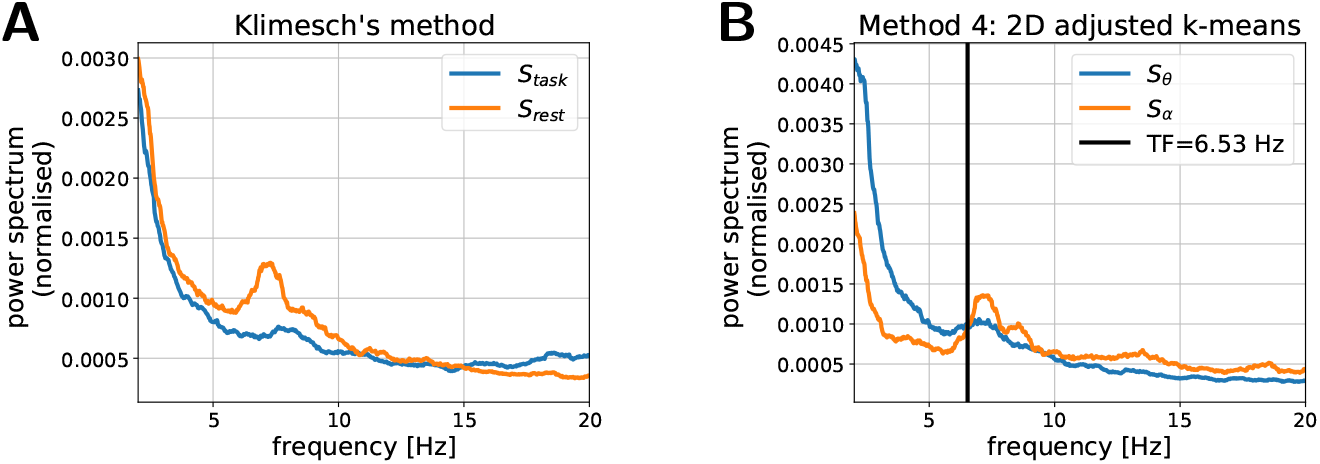
Example where Klimesch’s method cannot be applied because event-related, *S_task_*, and resting-state, *S_rest_*, spectral profiles do not intersect in a reasonable frequency range. (A) Results obtained with the Klimesch’s method. An event-related shift of the alpha–peak towards higher frequency can be seen in *S_task_*. (B) Results obtained with *transfreq* by using 2D adjusted k-means to compute the spectral profiles *S_θ_* and *S_α_*

*Transfreq* overcomes such limitations of Klimesch’s method, since it utilises just resting state recordings, and relies on the selection of specific channels that actually present the desired features, i.e. channels with a low (high) alpha and a high (low) theta activity for *G_θ_* (*G_α_*). Indeed, as shown in Figures 8 and 9, right panels, in both scenarios previously described *transfreq* estimated a reliable value for TF. More in general, a visual inspection of the results revealed that Klimesch’s method provided an untrustworthy value of TF for 27% of the EEG sessions of the open-source data set, while with *transfreq* only 6% of the results were unreliable.

### 3.4. Validation on the in-house data set

Figure 10 shows that the results obtained by applying *transfreq* on the inhouse data set are similar to those obtained on the open-source one. Specifically, *transfreq* generally returned higher estimates of TF with respect to Klimesch’s method. However, the absolute value of the difference between the values estimated with the two methods was below 1 Hz for 67% of the subject when 1D thresholding was applied, 58% with 1D mean-shift, 73% with 2D k-means, and 62% with 2D adjusted k-means.

**Figure 10:**
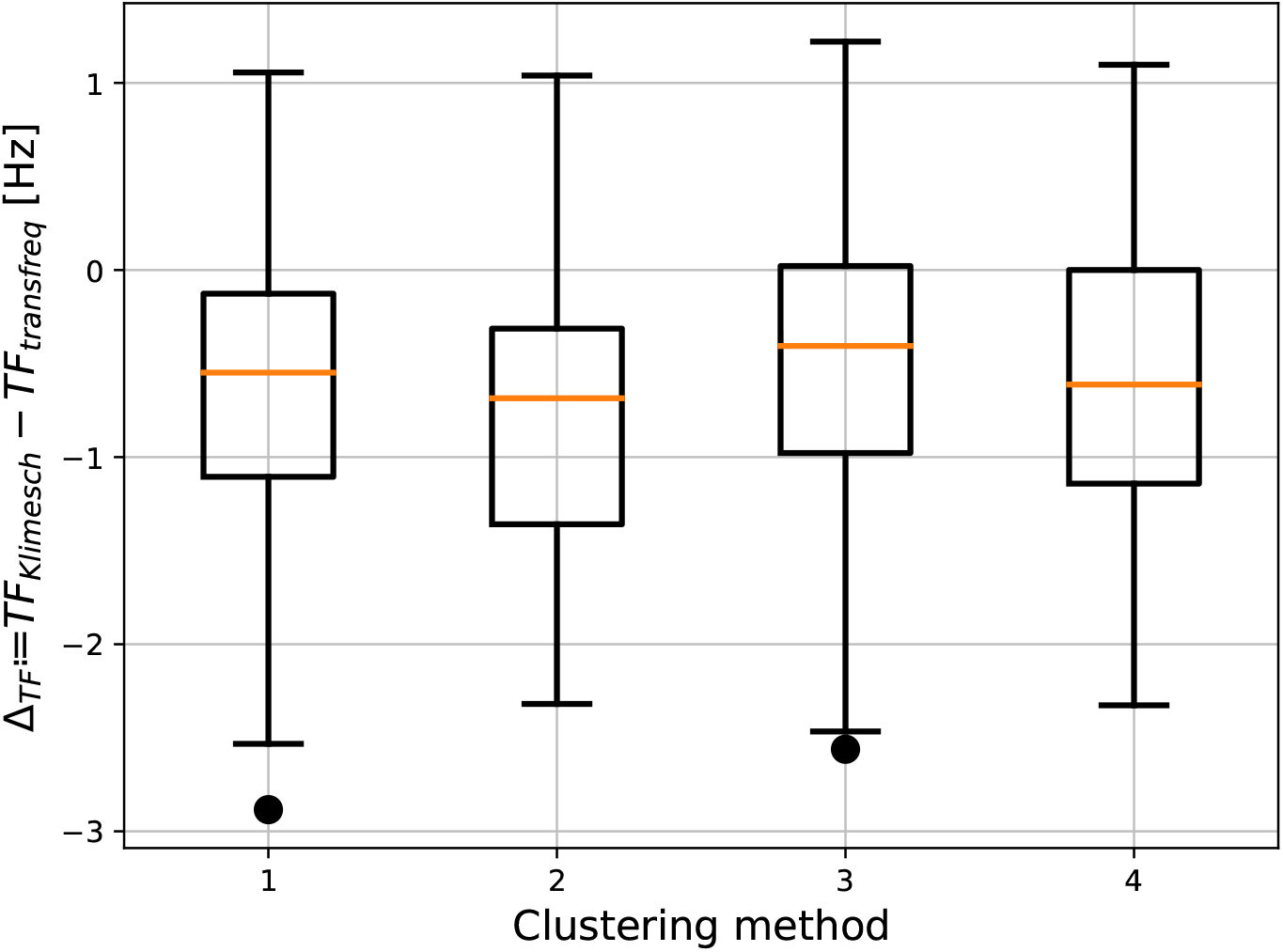
Difference between TFs estimated with Klimesch’s method (*TF_Kilimesch_*) and with *transfreq* (*TF_transfreq_*) over the in-house data set. As in Figure 7 each boxplot depicts the results obtained when a different clustering approach is used to define *G_θ_* and *G_α_*, namely: 1D thresholding (Method 1); 1D mean-shift (Method 2); 2D k-means (Method 3); and 2D adjusted k-means (Method 4).

### 3.5. Proportional bias in estimating TF

We performed a Bland-Altman analysis (Bland and Altman, 1986) to assess proportional bias in the estimates of TF. Figure 11 shows the analysis for the open-source (panel A) and the in-house data sets (panel B), computed on the TF values provided by *transfreq* with adjusted k-means. With the open source data set no proportional bias was present; to confirm this, we computed a regression line and the p-value (null hypothesis: slope equal to zero). Differently, the results with the in-house data set showed a statistically significant (*p* < 0.001) proportional bias. Specifically, Figure 11B shows that *transfreq* tends to overestimate TF at higher frequencies (> 8 Hz).

**Figure 11:**
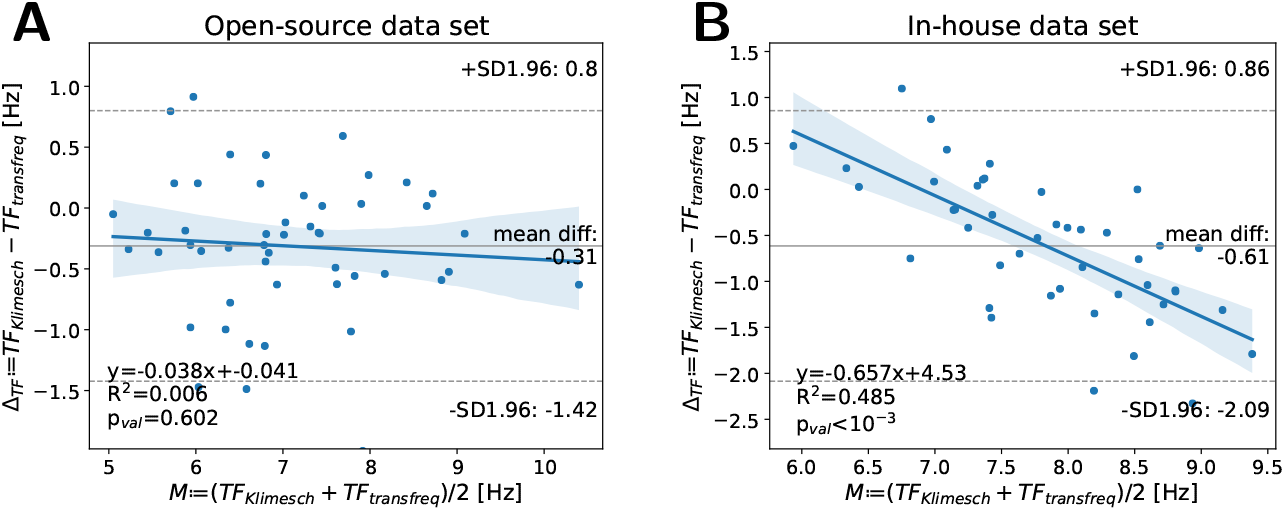
Bland-Altman plot between Klimesch’s method and transfreq with 2D adjusted k-means for the open-source (A) and the in-house (B) data sets. Grey plain and dotted lines show mean bias and corresponding 95% confidence limits, respectively. Proportional bias regression lines are depicted as blue lines, and the corresponding equations are embedded in the lower-left corner of each panel together with the coefficient of determination (*R*^2^) and the p-value (*p_val_*) computed testing the null hypothesis that the slope is equal to zero.

## 4. Discussion

A classic approach to compute the theta-to-alpha TF is that proposed by Klimesch and colleagues (Klimesch, 1999), which requires the power spectrum of two EEG time series, one recorded while the subject is resting and one while the subject is performing a task. However, in studies involving e.g. patients affected by neurodegenerative diseases, the subject may experiment difficulties in performing the required task and thus the corresponding event-related recording may imply difficult interpretation. On the contrary, *transfreq* uses only resting state data, which reduces the information at disposal but increases the scenario in which *transfreq* can be applied.

By comparing with the results obtained with the classic Klimesch’s method on two independent data sets, we demonstrated that *transfreq* returns reliable estimates of TF. Indeed, with the best combination of input parameters, the absolute value of the difference between the value of TF estimated with *transfreq* and with Klimesch’s method was below 1 Hz for 88% of the analysed data in the open-source data set, and for 73% for our in-house data set (throughout this paper Klimesch’s method was assumed as ground truth). The differences in the performance over the two data sets may be partially due to the noisier nature of the in-house data set. Moreover, a visual inspection of the estimated values of TF showed that the cases in which the spectral profiles, *S_θ_* and *S_α_*, obtained with *transfreq* intersected ambiguously were considerably less than the cases in which the hypothesis of Klimesch’s method on *S_task_* and *S_rest_* failed.

Among the four approaches implemented in *transfreq* to realise the clustering step, the adjusted k-means showed the best performances in the open-source data set while in the in-house data set the k-means algorithm performed the best. This is probably due to the fact that these algorithms realise a more accurate selection of the sensors within the two groups *G_θ_* and *G_α_*.

However, all four approaches tend to overestimate the value of TF with respect to Klimesch’s method. Specifically, the Band-Altman analysis for the in-house data set show that this behaviour seems to be more pronounced for higher values of TF (> 8 Hz). This difference between *transfreq* and Klimesch’s method is probably related to the fact that only resting-state data are used in *transfreq*; as a consequence also channels in *G_θ_* may present a fingerprint of the alpha activity.

The two data sets considered in this paper are EEG data. Future studies may be devoted to investigate a possible extension to MEG data.

Finally, future efforts will be devoted to better compare *transfreq* and the classic Klimesch’s method, especially in those scenarios where they return different estimates of TF. To this end, future studies will be devoted to correlate both Klimesch’s and *transfreq*’s results with clinical variables and biomarkers.

## 5. Conclusions

This paper introduces *transfreq*, an open-source Python tool for the computation of the individual transition frequency from theta to alpha band using only one resting-state EEG recording. The reliability of the obtained estimates was demonstrated by comparing the results of *transfreq* with those of the classic Klimesch’s method on two different data sets, one of which is publicly available.

